# HTLV-1 Splice Sites in Prevalent Gene Vectors Cause Splicing Perturbations in Transgenic Human Cells

**DOI:** 10.1101/2023.01.28.526022

**Authors:** Csaba Miskey, Sabrina Prommersberger, Katrin Mestermann, Michael Hudecek, Zoltán Ivics

## Abstract

The use of any semi-randomly integrating gene vector in a therapeutic setting is associated with genotoxic risks. The two major mechanisms of genotoxicity are disruption of a coding sequence (loss-of-function) or transcriptional upregulation of genes (gain-of-function) in the cellular genome where the genetic modifications are executed. A third, less widely recognized genotoxic risk stems from splice sites and polyadenylation sites within the vector sequences. These transcriptional elements may drive aberrant splicing and/or polyadenylation between transgene-contained and genomic sequences. A widely used promoter/enhance element present in gene vectors to ensure high transgene expression levels in mammalian cells is composed of a hybrid EF1α/HTLV-1 LTR, in which the retroviral LTR contains an intron. We assessed aberrant splicing initiated from the splice donor (SD) element present in the HTLV-1 LTR in CAR-T cells that had been engineered by either lentiviral vector (LV) or *Sleeping Beauty* (SB) transposon-mediated gene transfer. We establish that the vector-contained canonical SD site gives rise to aberrantly spliced RNA species and thereby can cause misexpression of host gene segments that are involved in various host cell functions. This, potentially genotoxic, effect could be abrogated by mutating or completely eliminating the SD (or the entire intron) from the HTLV-1 LTR segment. CAR-T cells generated by the modified vectors are equally potent in efficiency of CAR-T cell manufacturing and in functionality. The simple genetic modifications that we describe here affecting vector design therefore enhance genomic safety while maintaining efficacy of gene-modified therapeutic cells.

## INTRODUCTION

The currently available integrating gene vector systems (either virus- or transposon-based) all insert their genetic cargo into the genome in a semi-random fashion. The random nature of integration can result in two major issues, both of which are especially problematic for therapeutic applications. The first problem, termed position effect, is that expression of the transgene can be influenced by its position in the genome. This may lead to unpredictable levels of expression of the integrated genetic cargo, which in turn leads to unpredictable therapeutic efficiency and side effects^1^. The other problem, genotoxicity, is that insertion of a transgene can result in the disruption or dysregulation of genomic elements, and these mutagenic events are especially problematic if proto-oncogenes or tumor suppressor genes are affected, because deregulation of these genes can result in uncontrolled proliferation of cells and the development of tumors *in vivo* (e.g., ^2^). Such risk is especially pronounced with gammaretroviral vectors based on the Murine Leukemia Virus (MLV) that preferentially integrate into transcriptional regulatory elements of active genes (e.g., ^3^; in fact, severe adverse events associated with vector integration have been observed in clinical trials for SCID-X1 (e.g., ^2^) and for other monogenic diseases affecting the hematopoietic system. HIV-derived lentiviral vectors (LVs) are also potential mutagens due to their biased insertion into transcription units^4^.

We have previously undertaken a comparative study addressing target site selection properties of the *Sleeping Beauty* (SB) and *piggyBac* transposons as well as MLV-derived gammaretroviral and HIV-derived lentiviral vector (LV) systems in primary human CD4^+^ T cells. Our bioinformatic analyses included mapping against the T cell genome with respect to proximity to genes, transcriptional start sites (TSSs), CpG islands, DNaseI hypersensitive sites, chromatin marks and transcriptional status of genes. The SB transposon displayed the least deviation from random with respect to genome-wide distribution: no apparent bias was seen for either heterochromatin marks or euchromatin marks and only a weak correlation with transcriptional status of targeted genes was detected ^5^. Our compiled datasets also allowed us to rank these vector systems with respect to their projected relative “safety” based on the frequencies of integration into genomic safe harbors (GSHs) that were previously proposed to satisfy five criteria: (i) distance of at least 50 kb from the 5ʹ-end of any gene, (ii) distance of at least 300 kb from any cancer-related gene, (iii) distance of at least 300 kb from any microRNA (miRNA), (iv) location outside a transcription unit and (v) location outside ultraconserved regions of the human genome^6^. Our analyses collectively established a favorable integration profile of the SB transposon, suggesting that SB might be safer for therapeutic gene delivery than the integrating viral vectors that are currently used in clinical trials.

The available data indicate that transcriptional upregulation of proto-oncogenes by transgene-associated, internal enhancers and promoters is a major mechanism of insertional oncogenesis in gene therapy clinical trials. However, integrating viruses, and thus the gene vector systems that have been developed from them, also tend to contain polyadenylation (polyA) signals and splice sites. Once integrated in or close to genes, these sequences could potentially interfere with the initiation and processing of cellular mRNAs, create truncated proteins with stimulatory or inhibitory functions, or lead to monoallelic inactivation of tumor suppressors. For example, proto-oncogene activation may involve the generation of chimeric transcripts originating from the interaction of promoter elements or splice sites contained in the integrating genetic element with the cellular transcriptional unit targeted by integration^7,8^. Chimeric fusion transcripts comprising vector sequences and cellular mRNAs can be generated either by read-through transcription starting from virus/vector sequences and proceeding into the flanking cellular genes or vice versa^7–10^. For example, earlier studies have identified provirus-dependent host gene perturbations as drivers of leukemogenesis in human T-cell leukemia virus type-1 (HTLV-1) infected cells^11,12^. These studies showed that splice donor (SD) and acceptor (SA) sequences present in the retroviral LTRs are involved in non-physiological *cis*-splicing events, which take place between the viral and host gene sequences. It was found that in leukemia patients, the aberrant splicing events tend to occur with genes that are known cancer drivers; hence, these perturbations play an important role in tumorigenesis. Similar, the long terminal repeats (LTRs) of lentiviral vectors (LVs) contain both canonical and cryptic polyA signals in both orientations^13^, and can therefore elicit premature termination of gene transcription. These polyA sites, however, are used only in a context-dependent manner and when specific requirements are met^14^. Moreover, viral-cellular fusion transcripts terminating at cryptic polyA sites located in the host cellular genome were found in keratinocytes transduced with LVs^15^. In addition, both in a β-thalassemia clinical gene therapy trial as well as in a murine model aberrantly spliced products of endogenous transcripts fused to vector sequences led to either proto-oncogene upregulation or haploinsufficiency of a tumor suppressor gene, respectively^10,16–18^. Noteworthy, in most cases engagement of splice sites within the LV leader sequence contributed to abnormally spliced transcripts^16,17^. However, recent reports have revealed that cryptic splice sites are also engaged in splicing leading to aberrant cellular exon–vector fusion transcripts^16–18^. In sum, a complex interplay among the presence, relative strength, position, and distance of promoters; splice sites and polyA signals is ultimately responsible for the production of specific aberrant mRNAs. Thus, there is emerging evidence that the potential of inducing aberrant transcripts might constitute a previously underappreciated genotoxicity factor for gene therapy vectors.

Intriguingly, the elongation factor alpha 1 (EF1α) promoter-enhancer segment of a commonly used gene vector construct contains the R segment and part of the U5 sequence (R-U5’) of the HTLV-1 LTR, including SD and SA sequences previously shown to be engaged in ectopic splicing in HTLV-1 induced leukemia patients^19^. To assess the transcriptional safety of the transgene construct, we measured the involvement of the retroviral splice sites in the splicing of the transcripts originating from this hybrid transgene promoter, both in the context of SB transposon and LV vectors. Variants of the SB-based vectors were engineered with and without a doublet of the SV40 polyA signal^20^, a base substitution or deletion mutagenizing the SD or complete removal of the intron from the HTLV-1 LTR segment. These vectors were then used for the generation of CAR-T cells, which were then used to validate CAR expression, CAR-T cell function and to both qualitatively and quantitatively annotate and measure splicing events between vector sequences and endogenous human genes. We conclude that full deletion of the intron from the HTLV-1 segment of the transgene promoter as well as inclusion of strong polyadenylation in the transgene cassette almost completely abolish aberrant, vector-genome splicing events. These observations not only inform us with respect to future vector designs but significantly improve genomic safety of therapeutic cell products engineered with semi-randomly integrating gene vectors.

## RESULTS

### Inefficient transgene polyadenylation and active HTLV-1 LTR-derived spice sites in gene vectors cause host gene misexpression in transgenic CAR-T cells

To assess the transcriptional safety of the transgene construct, we measured the involvement of the retroviral splice site in the splicing of the transcripts originating from the transgene promoter (**Figure 1A**).

**Figure 1.**
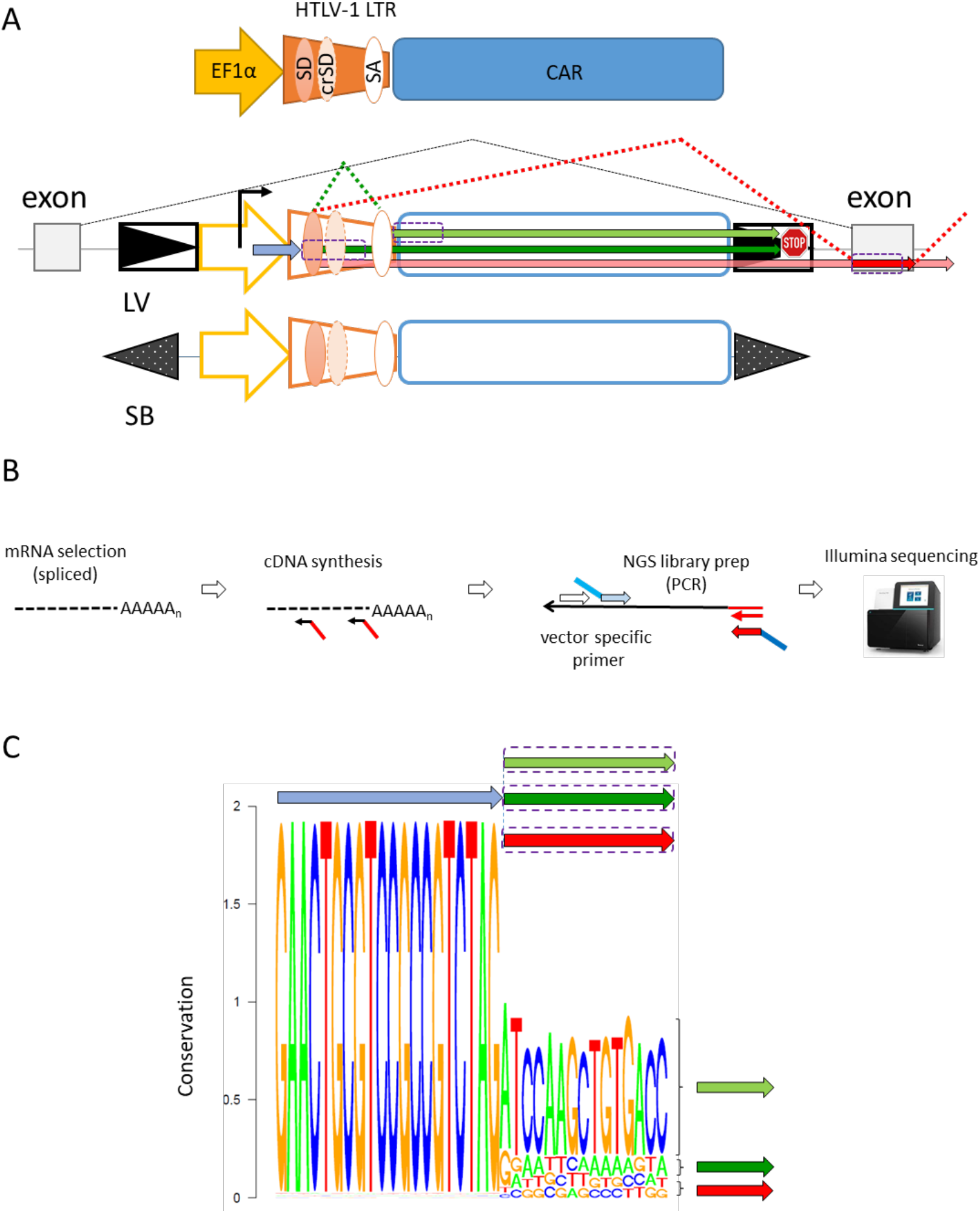
HTLV-1 LTR fragment in the promoter/enhancer region of the gene vector can generate splice products with sequences within and outside of the transgene. **(A) Schematic of the transgene with the position of the HTLV-1 LTR, the splice sites, and the most abundant RNA products.** Transcripts (colored arrows) originating from integrated SB and LV vectors may go through splicing within the LTR sequences. Those transcripts, which are not polyadenylated at the 3’ end of the vector, can be engaged with splicing with host sequences downstream of the vector. Canonical and cryptic splice donor sites (SD and cSD, respectively) are shown with orange ovals, the splice acceptor site (SA) is shown in white. The framed triangles stand for the LTR of the LV vector; dotted ones represent the inverted repeats of the SB transposon. The stop sign stands for a polyadenylation signal sequence. **(B) Workflow of the targeted RNA-Seq assay for the detection of splice products.** After cDNA synthesis performed on polyadenylated RNA using random hexamers, nested PCR, specific for the HTLV-1 LTR is used to amplify cDNA molecules spanning the region of interest. The position of the primer (in blue) specific for the LTR sequence upstream to the SD site is shown in (A). **(C) The splicing events are classified by the sequence composition of reads following the splice site.** The sequence logo shows the most abundant splice products by their sequence conservation (*y*-axis, maximum conservation is log_2_ 4=2). The green and red arrows show the predominant splice products, the colors correspond to the major transcripts depicted in (A).

We generated CAR T cells using an LV- and an SB system-based vector expressing a ROR1-specific CAR from the hybrid EF1α/HTLV-1 LTR promoter-enhancer element. CD4^+^ and CD8^+^ T cells were isolated from leukocyte reduction chamber systems and subsequent magnetic bead separation; and were activated with Transact. After one day of activation, T cells were transduced with 3 MOI of LV or, after two days, transfected with SB100X transposase-encoding mRNA and CAR-encoding minicircle (MC) DNA. T cells were enriched for CAR expression and expanded by stimulation with irradiated feeder cells and OKT3 antibody.

We devised a high-throughput, targeted RNA sequencing (RNA-Seq) experiment to selectively amplify transcripts, which are driven by the composite EF1α/HTLV-1 LTR promoter-enhancer segment (**Figure 1B**). RNA species containing the HTLV1 LTR sequences upstream to the described SD site can be quantified based on the sequence compositions of the reads following the canonical SD dinucleotide signature. We identified the major types of RNA species and the causal splicing events, with the potential engagement of the known SD sequence (**Figure 1C**). The three major transcript types were: (i) transcripts, which were not subjected to splicing at the SD, but continued with the downstream vector sequence; (ii) RNA species, deriving from splicing between the SD and a splice acceptor (SA) site situated downstream in the HTLV-1 LTR segment; and (iii) transcripts, which underwent splicing, but the SAs that had been engaged in splicing were other than of vector origin. Since the sequences of the first two groups are both unique, the above three groups are distinguishable on the sequence logo made of the reads following the SD site (**Figure 1C**). We also identified a small number of reads (<0.01%), which indicated the adoption of a so far unknown cryptic SD site, 15 nucleotides downstream to the canonical one in the LTR segment (**Figure 1 A**). The results above suggested that active SD sites within the HTLV-1 LTR segment of integrated gene vectors give rise to aberrant transcripts in transgenic CAR T cells.

Next, we aimed at identifying the genomic locations of those reads containing non-vector sequences downstream of the vector SD site by mapping these to the human genome. We considered any uniquely mapped position as a valid one for downstream analysis, if it was supported by at least ten independent reads. Consequently, the numbers of the genomic loci considered for downstream investigations are likely underestimations. Retrieving the sequences upstream to the mapped positions revealed that these loci are indeed canonical SA sequences (by their nucleotide composition) and the splicing events between the vector and the genomic splice sites followed the established rules of vertebrate splicing (**Figure 2A**).

**Figure 2.**
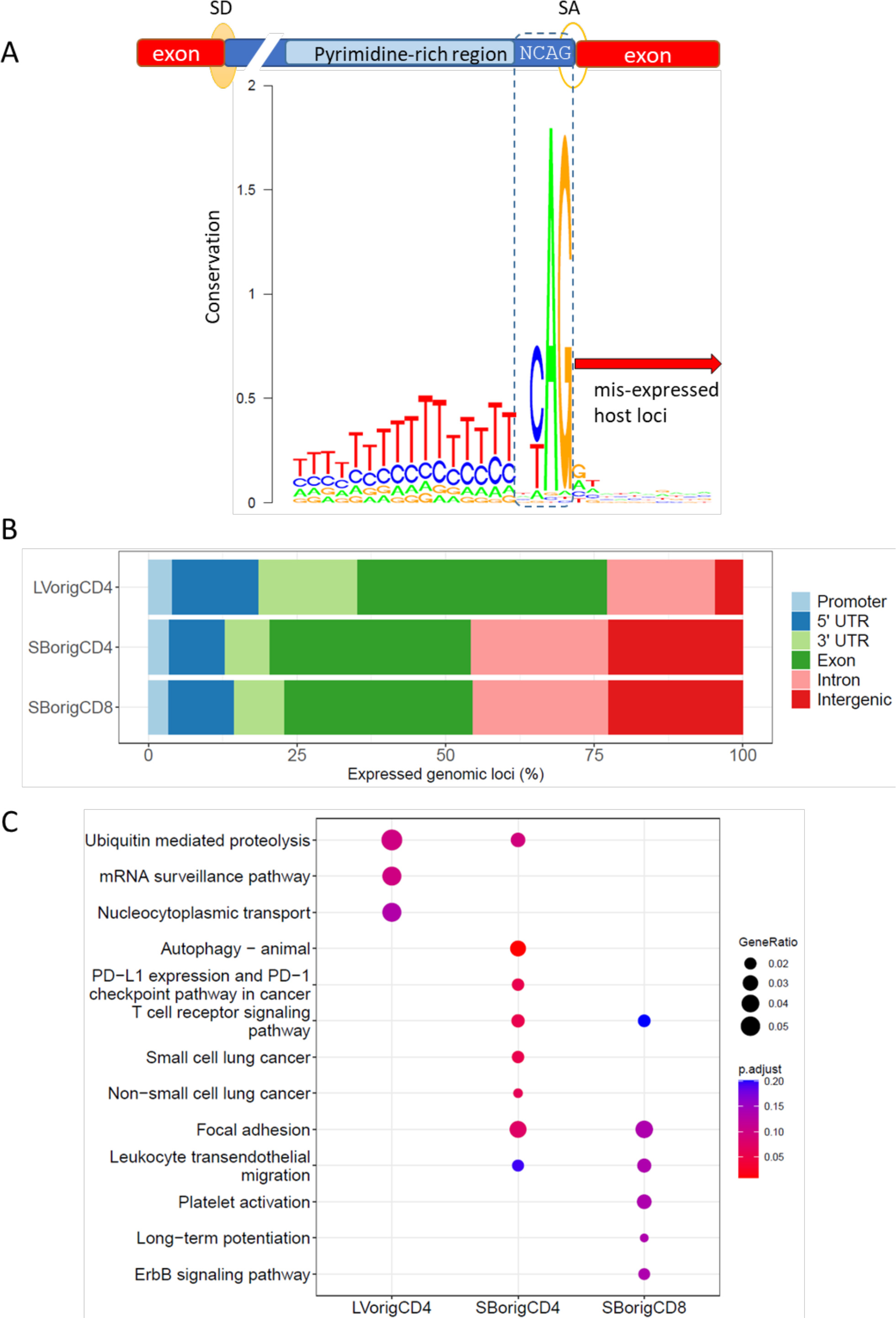
Characterization of the ectopically expressed host sequences, which arise via *cis*-perturbations. **(A) Downstream host segments are included in the vector-initiated transcripts by canonical splicing.** Above, conserved sequence composition of the SA sites in pre-mRNA molecules of vertebrates. Below, the sequence logo shows the base composition of human genomic loci immediately upstream to the spliced segments, present in the mRNAs from cells transgenic for the LV vector. The conserved intronic sequences at the 3’ splice sites are framed. SD and SA stand for splice donor and acceptor loci, respectively. **(B) Most of the misexpressed downstream genomic segments are known human exons.** Genomic distribution of host sequences involved in splicing with the transgene RNA are shown. The percentage representation in gene-related categories (on the left) are shown on the *x*-axis. **(C) Transcriptional perturbations can affect host cell-specific functions**. Listed are the most significantly enriched pathways obtained by KEGG pathway enrichment analysis. The overrepresentation analysis took account of those genes that were involved in misexpression *via* vector RNA-genome splicing.

We then overlapped the genomic coordinates of the vector-host junctions with the positions of gene-related segments of the human genome. As expected, most of the misexpressed segments were fragments of genes, and intergenic regions were underrepresented, particularly if the expression was initiated from within an LV vector (LV: 4.6%, SB: 22.2% and 22.5% in CD4 and CD8 positive T cells, respectively; **Figure 2B**), consistent with the characteristic difference of LV and SB vectors to target expressed genes for integration^5^. Most of the misexpressed sequences were known exons (LV: 32.7%, SB: 29.5 and 28.2%). We found that a larger portion (LV 22.6% vs. SB 12.5% and 12.2%) of the overexpressed loci overlapped with gene promoters, if the latter category was defined as windows 3kb upstream to transcriptional start sites (TSS).

To determine what biological processes and pathways are affected by the misexpression, we performed a gene enrichment analysis based on the list of genes, whose segments were involved in chimeric splicing. The analysis, for which we harnessed the collection of the Kyoto Encyclopedia of Genes and Genomes (KEGG, Ref.^21^), identified several significantly (FDR>0.1) enriched pathways, including processes of T cell-related functions, including trans-endothelial migration and T cell receptor signaling (**Figure 2C**). In sum, the results above suggested that the primary transcripts originating from the promoter segment of the CAR transgene are not polyadenylated efficiently within the transgene vectors in T cells. Consequently, the integrated strong promoter/enhancer sequences in the combination with a canonical SD site could give rise to aberrantly spliced RNA species and thereby can cause misexpression of host gene segments involved in various host cell functions.

### Sequence modifications of the SB CAR T gene vector abolish splicing initiated from the HTLV1 LTR

We assumed that the decisive factors in the emergence of the abnormal RNA species described above are inefficient polyadenylation of the primary transcripts, and the presence of a strong SD sequence close to the 5’ end of the primary RNA. Therefore, we created mutant derivatives of the SB gene vector harboring polyA sites at the 3’-end of the transgene construct, and mutation/deletion in the 5’ regions active in splicing (**Figure 3A**). Three new constructs were derived: in the construct (1) “SBmutSD” the SD sequence was mutated by changing the GT nucleotides to TG, in (2) “SBdelSD” the GT nucleotides of the SD sequence were removed completely and in (3) “SBdellntr” the LTR sequence was truncated by the nucleotides reaching from SD to SA (remaining LTR sequence counts 123 bp). All three vectors contained two SV40 poly A signal sequences (each 122 bp) after the stop codon of the transgene, following previous observations^20^. Of note, the promoter regions and the CAR coding sequences are identical in the “SB orig” and “LV orig” vectors.

**Figure 3.**
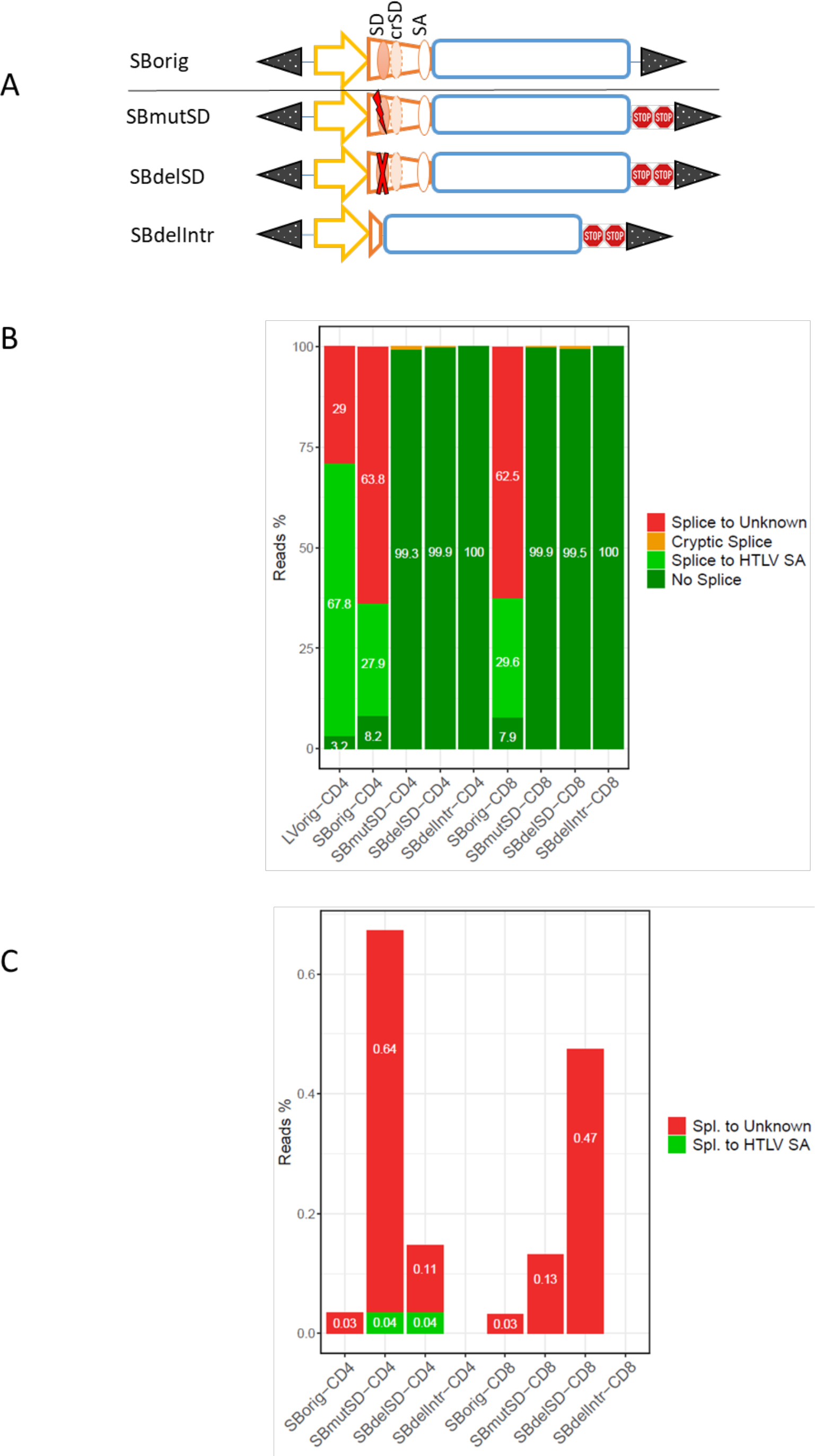
Vector modifications suppress splicing initiated from the HTLV-1 LTR of transgene constructs. **(A) Schematic structure of the original and the modified SB gene vectors.** Nucleotide substitutions (SBmut), or deletion were applied on the canonical SD site in the HTLV-1 LTR. The complete intron of the LTR (including all splice sites) was removed to obtain the SBdelInt derivative. The stop signals represent SV40 late polyA signals. **(B) Quantification of splice products from integrated SB and LV transgenes in CD4+ or CD8+ CAR-T cells**. Percentage representations of all transcript classes (explained in Figure 1) are shown on the *x*-axis. **(C) Disruption of the canonical SD site increases the frequency of splicing initiated at a downstream cryptic SD signature.** Shown are the percentages of reads, in which the cryptic SD site of the HTLV-1 LTR was used for splicing with the vector SA, or with downstream genomic sequences.

The original and the modified minicircle (MC) vectors were used to generate CAR-T cells. Polyadenylated RNA samples from cells transgenic for the original (SB and LV) and the modified SB vectors were used to quantify RNA species containing sequences adjoining the splice sites (**Figure 3B**). The majority (67.8%) of the sequences showed splicing between the canonical SD and SA sequences of the viral LTR in CD4 positive CAR T cells harboring the LV vectors. We found that 29% of the LV vector derived transcripts continued with sequences other than the expected LTR sequences after the SD site, or than the sequence deriving from splicing within the LTR. Strikingly, we found that the majority of the sequences following the SD locus was of non-vector origin in CD4^+^ and CD8^+^ CAR-T cells transgenic for the original SB CAR transposon (63.8% and 62.5%, respectively). The second most abundant reads were derived following splicing through the canonical SD and SA sequences of the viral LTR of the promoter (27.9% and 29.6% in CD4^+^ and CD8^+^ cells, respectively). On the contrary, the overwhelming majority (>99.3%) of the transgene-derived RNA continued with the expected vector LTR sequence both in CD4^+^ and CD8^+^ CAR-T cells generated by the modified SB vectors. The frequency of splicing using the cryptic SD (cSD) site was generally much lower (<1%), compared to the original SB, in all the conditions and showed an increased occurrence in vector mutants, in which the canonical SD site was either mutated or deleted (**Figure 3C**). Interestingly, splicing between the cryptic SD and the cognate SA within the vector was inefficient and the majority of the cryptic RNA fusions made use of downstream SAs, outside of the vector-encoded RNA. These results show that the removal of the critical nucleotides for splicing by either nucleotide substitution or deletion abolish the possibility for splicing at the canonical SD site. This, however, results in an increased frequency of cryptic splicing events. As expected, deleting all of the LTR intron, including splice signatures, eliminates the possibility for splicing at these vector sequences.

### The SB100X vector system shows safer genomic insertion profile than the HIV lentivirus-based gene vector

In order to profile the genomic insertion of the gene vectors we performed targeted DNA sequencing of genomic segments bordering the LV and SB integrations in CAR-T cells^22^. The insertion sites were determined by stringent mapping of the high throughput sequencing reads to the human reference genome (hg38). Both of the vectors exhibited the well-known preferences for the nucleotide composition of the insertion sites (**Supplementary Figure 1**). In accordance with earlier results^23^, we found that the LV insertions are more biased towards genes than those of the SB system (**Figure 4A**). In fact, only 7.8% of the LV insertions took place outside of genes; while 32.2% of the SB insertion sites were intergenic in CD4^+^ CAR-T cells. The two vectors showed similar frequencies of integration in the close vicinity of TSSs (**Figure 4B**). However, the LV-mediated insertions were biased towards regions downstream of the loci of transcriptional initiation. Indeed, studying the distribution of the insertions in and around genes recapped the known preference of HIV and of LV to insert inside gene bodies (**Figure 4C**). On the contrary, SB insertions show no frequency biases within or at the boundaries of genes. In general, we found that the introduced modifications in the transcriptionally active segments of the transgenes did not invoke significant changes in the insertion site distribution of the SB vector. In summary, the results of our insertion site profiling studies are in good agreement with previous findings, showing an overall safer genomic integration profile of SB than of LV gene vectors.

**Figure 4.**
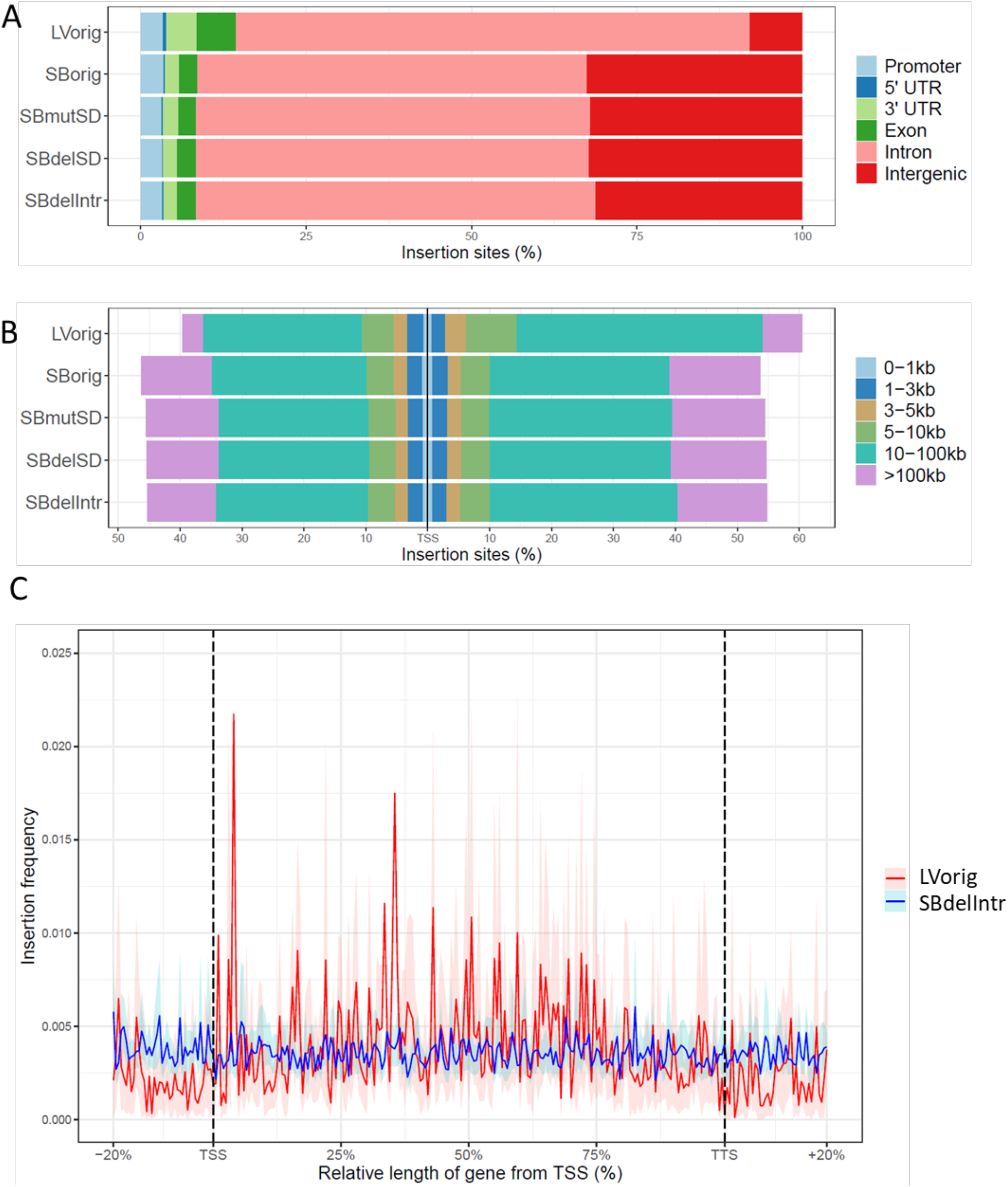
Insertion site distributions of the original (LV and SB) and the modified SB gene vectors in the genome of human CD4+ T cells. **(A) Representation of integrations in gene-related genomic segments.** The coordinates of the categories (on the left) were retrieved from the RefSeq database (hg38). Promoters were defined as regions 3kb upstream from the transcriptional start sites (TSS) of known genes. **(B) Distribution of insertions around transcriptional start sites.** The frequencies of the insertions were determined in genomic windows of increasing widths around TSSs (on the left). **(C) Localization of insertions in gene bodies.** The coordinates of all known genes were binned to 200 segments and the insertions were counted in each bin. TSSs and transcriptional termination sites (TTS) correspond to 0% and 100% of gene length, respectively. The light blue and red stripes show the confidence interval of 99%)

### Sequence modifications in MC construct have no impact on T cell transfection efficiency

Next, we tested if the changes in promoter and polyA signaling regions have any impact on the efficiency of CAR-T cell generation. T cells were transfected either with the SBoriginal or the SBdellntr MC and expanded for 10 days. Afterwards, transfection efficiency was analyzed by flow cytometry. We found no significant difference in the expansion rates of CD4^+^ and CD8^+^ T cells after using one of the two MCs (data not shown). Further, we found that transfection with the SBdellntr MC does not lead to any difference in transgene transfection rates with regards to percentage and mean fluorescent intensity compared with the transfection of SBorig MC (**Figure 5A**). Importantly, there were also no differences in the T cell phenotype with regard to subset and activation/exhaustion markers (**Figure 5B**).

**Figure 5.**
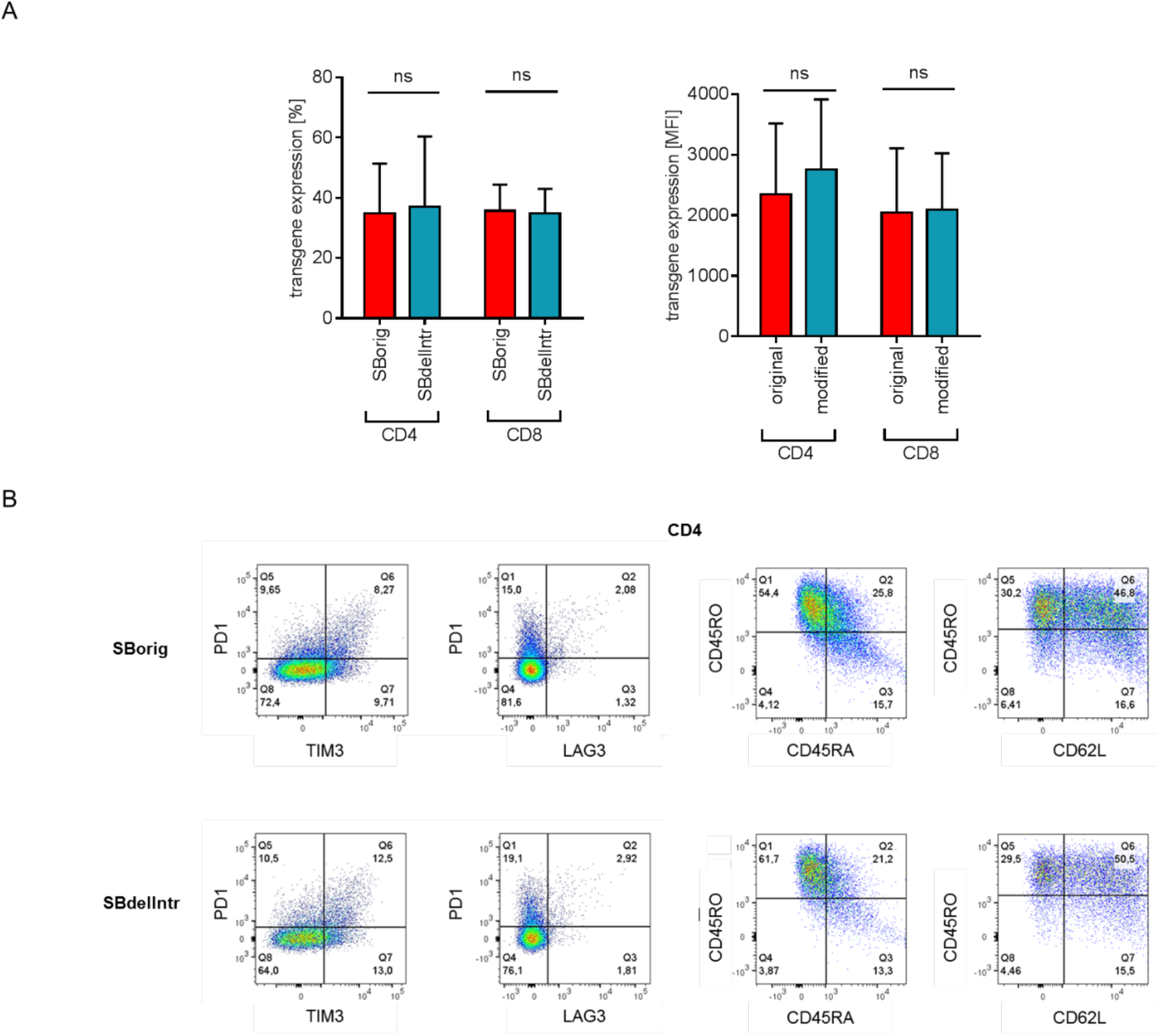
Transgene expression and T cell phenotype after transfection with both MC variants. CD4 and CD8 CAR-T cells were generated using the SBorig or the SBdellntr MC version. **(A)** After harvest on day 12, transgene expression was measured by flow cytometry (n=6 CD4, n=7 CD8, mean +SD). **(B)** Further, expression of T cell activation and subset markers on CAR-expressing T cells was measured by flow cytometry.

### CAR-T cell functionality is not impaired by modifications of the transgene sequence

Finally, we analyzed if the transgene design has any impact on the functionality of CAR-T cells. Therefore, CAR-T cells were generated using the SBorig or the SBdellntr MCs, and co-incubated with target antigen-negative K562 control cells, with K562 cells ectopically expressing the target antigen or with the breast cancer cell line MDA which endogenously expresses the target antigen. CAR-T cells eradicated both target antigen-expressing cell lines while sparing control cells. There was no difference in killing efficiency detectable when the T cells had been transfected with the one or other MC (**Figure 6A**). Same was true when analyzing cytokines secreted by CAR-T cells. There was no significant difference of IL-2 and IFN-y secretion levels by both types of CAR-T cells after stimulation with target antigen-expressing cells (**Figure 6B**). We finally compared the capability of CAR-T cells transfected with the SBorig or SBdellntr MC vector to proliferate after antigen encounter. We found similar proliferation profiles for both types of T cells (**Figure 6C**). To summarize, the usage of the optimized MC SBdellntr for gene transfer did not induce any difference in efficiency of CAR-T cell manufacturing or in the functionality of the CAR-T cells.

**Figure 6.**
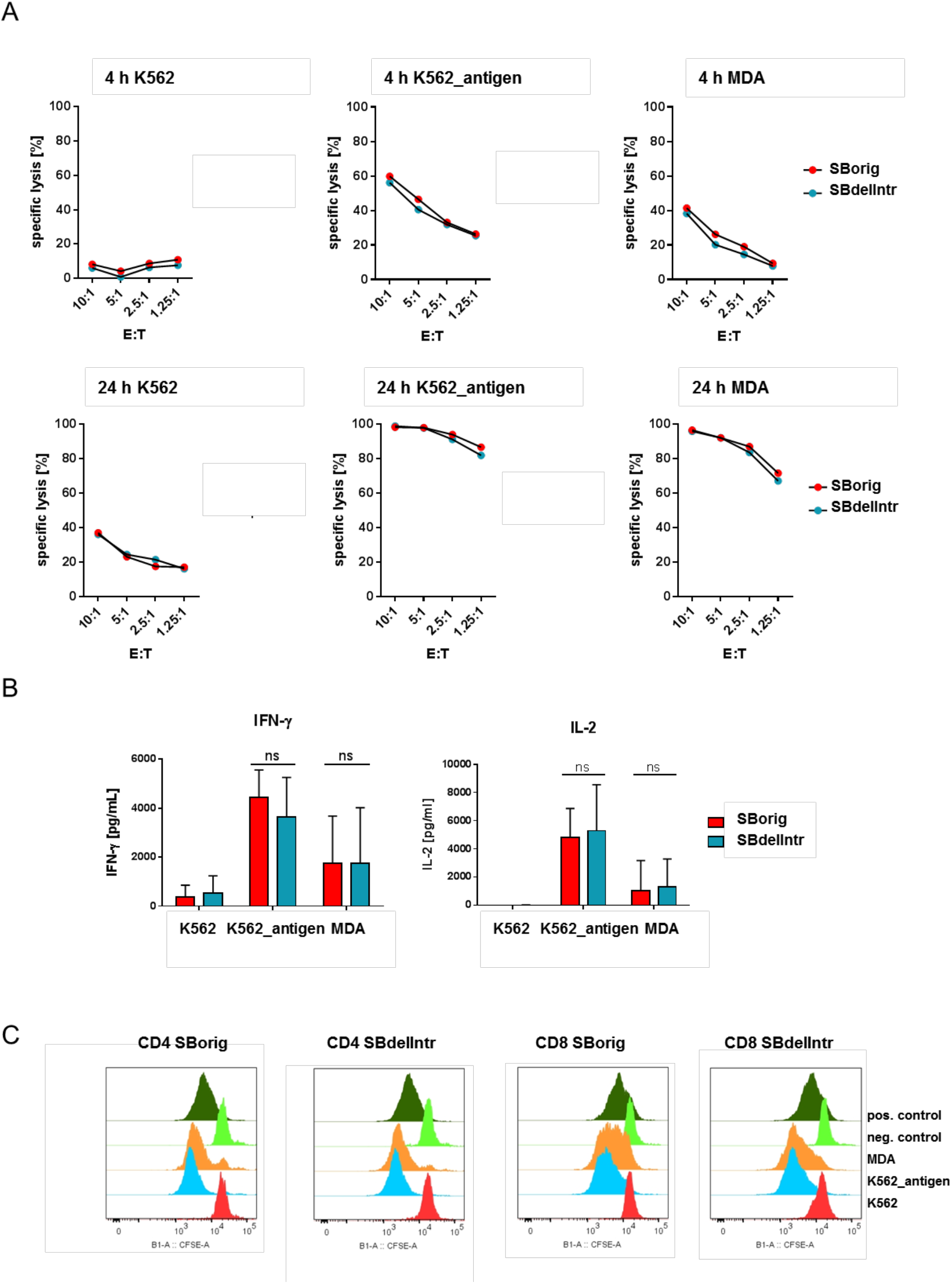
Functionality of CAR-T cells either generated with the SBorig or the SBdellntr MC. CAR-T cells were generated either using the SBorig or the SBdellntr MC for nucleofection. **(A) CAR-T cells were tested in luciferin-based cytotoxic assays for their lytic capacity.** Antigen-expressing target cells (K562_antigen, MDA) or antigen-negative cells (K562) were used as targets. Specific lysis was analyzed after 4 and 24 hours. Experiment was performed in technical triplicates. Graphs show mean values from n=3 different donors. **(B) Cytokine secretion by CAR-T cells after 24 h of stimulation with antigen-expressing target cells or control cells was measured by ELISA.** Experiment was performed in technical triplicates. Graphs show mean values from n=3 different donors. **(C) CAR-T cells were labeled with CFSE and stimulated for 72h with antigen-expressing target cells or control cells.** Representative data set of n=4 experiments.

## DISCUSSION

In this study, we described how HTLV-1 LTR-derived sequences, present in common gene vector constructs, provoke transcriptional perturbations and host gene misexpression in transgenic cells. We showed that a strong SD signal in the viral LTR-derived enhancer/promoter fragment is mainly responsible for driving activating transcript fusions with gene fragments downstream to the transgenes. We found that the majority of the misexpressed sequences are known exons of host genes, which are included in the transgene-derived transcripts by canonical splicing. We showed by gene enrichment analysis that the affected genes are often involved in host cell related functions.

The two prerequisites behind the emergence of the aberrant splicing products are the presence of the SD site in the 5’-end of the transcripts and inefficient polyadenylation of the RNA molecules. Therefore, we modified the vector constructs by mutating and deleting the splicing-active segments of the vector and by including duplicated SV40 polyA sites in to 3’-end of the gene vectors. Nucleotide substitutions or deletions of the conserved dinucleotides of the canonical SD site increased the frequency of splicing at a cryptic and so far undescribed SD site within the HTLV-1 LTR fragment of the promoter. By removing the entire intron and the splice site from the promoter construct, the possibility of splicing was abrogated from the vectors.

We measured significant differences between the intragenic representations of the misexpressed genomic loci if the cells were transgenic for the LV *vs*. the SB vectors (95.4% and 77.5%, respectively, **Figure 2B**). Our genome-wide insertion site profiling results indicated that this phenomenon is due to the differences in the overall insertion site distributions of the two vector systems: the LV insertions being biased towards genic segments, while SB transposition lacks this tendency.

Finally, we have shown that the genetic changes we introduced into the transgene cassettes did not have any measurable impact on the biological properties of the resulting CAR-T cell products. The data therefore argue that our vector modifications do result in significant enhancement of genomic safety in therapeutic cell engineering while maintaining potency and efficacy of the gene-modified cell product.

## MATERIALS AND METHODS

### High-throughput targeted sequencing of the vector HTLV LTR-derived transcripts and the insertion sites of the SB and LV vectors

Total RNA samples from CART cells were isolated with the DirecZOL RNA Miniprep (Zymo Research). One μg of total RNA was used to select polyadenylated RNA using the NEBNext® Poly(A) mRNA Magnetic Isolation Module (New England Biolabs, NEB). Half of the mRNA per sample was used for cDNA synthesis using the Superscript IV transcriptase (Thermo Fisher Scientific) following the recommendation of the manufacturer at 45 °C for 30 min with the NNSR-2 primer (see table of primers below). The reactions were supplemented with RNaseH (NEB) and incubated at 30 minutes at 37 °C. Following magnetic bead purification one third of the cDNA per sample was subjected to PCR amplifications with the primers HTLV-1 and PE-2nd-GA using the NEBNext® Ultra™ II Q5® Master Mix (NEB) using the cycling program: 98 °C 30s; 15 cycles of 98 °C 10s, 70 °C 40s; 15 cycles of 98 °C 10s, 57 °C 20s, 72 °C 40s. A nested PCR round was harnessed to append the library with sample specific barcodes and Illumina adapters with the primers NNSR-indN and HTLV-sd-bcN, where N stands for the number of the Illumina TruSeq adapter sequences. The PCR program was 98 °C 30s;15 cycles of 98 °C 10s, 70 °C 40s. The PCR products were separated in an agarose gel and the smears of 250-500 bp were isolated and sequenced with an Illumina NextSeq 550 instrument using a single end 86-nucleotide setting.

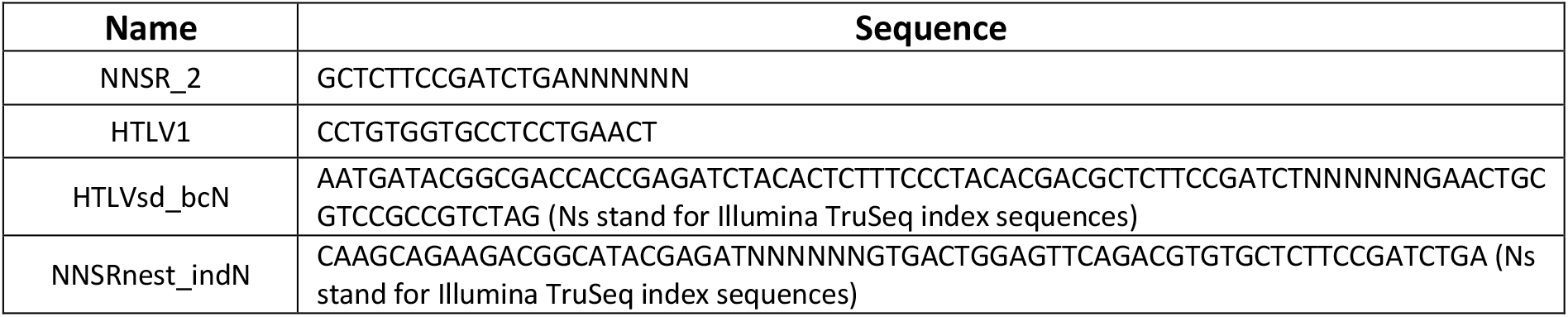

### Quantification of the HTLV LTR-derived transcripts, analysis of misexpressed genomic segments and the vector insertion sites

We used *fastp^24^* for the quality-, adapter-, and minimum length-trimming of the raw reads. Pre-filtering of the reads for the presence of the LTR specific sequences and the downstream analysis was done using the R environment https://www.R-project.org/. Reads were filtered for the presence of the HTLV-1 specific primer sequence. The splicing products within the vector using the canonical or the cryptic SA sites and non-spliced transcripts were identified by their sequence composition and quantified. Sequences containing non-vector sequences after the SD sites were designated as “spliced to unknown” and were further analyzed as follows. We used *STAR^25^* to map the unknown segments of the reads with a minimum length requirement of 56 to the human genome assembly (hg38). The mapping output was filtered by MAPQ>10 and the mapping positions were tested for the presence of a sequence similar to the vector specific primer, in order to filter out possible (but rare) mispriming events. Also, any mapped genomic position had to be supported by at least 10 independent reads. We used *ChIPseeker* to annotate the positions of the spliced sequences in the human genome and used *clusterProfiler^26^* for gene enrichment analysis in the pathways of the Kyoto Encyclopedia of Genes and Genomes^21^. Targeted DNA-sequencing of the insertion sites, their quantification and annotation has been described^23,27^. In addition to the tools described above we used *ChIPseeker^28^* to annotate the insertion sites in gene related segments of the human genome.

### Generation of CAR-T cells

We generated CAR-T cells using an LV and the SB100X (SB) system-based CAR vector. Therefore, CD4^+^ and CD8^+^ T cells were isolated from leukocyte reduction chamber systems by gradient centrifugation over Pancoll separating solution (Pan-Biotech, Aidenbach, Germany) and subsequent magnetic bead separation (Miltenyi, Bergisch Gladbach, Germany) and activated in T cell medium (1640 RPMI with HEPES + Glutamax, 10% human serum, 1% Glutamax, 1% penicillin/streptomycin, 0.1% β2-mercaptoethanol) supplemented with Transact (Miltenyi, Bergisch Gladbach, Germany) and human IL2. After one day of activation, T cells were transduced with 3 MOI of lentivirus or, after two days, transfected with SB100X-encoding mRNA and CAR-encoding minicircle (MC) using a Maxcyte (Gaithersburg, US) electroporater device and program “Expanded T cell 3”. T cells were expanded in T cell medium with 50 U/ml IL-2. At day 13, T cells were enriched for CAR expression and expanded by stimulation with irradiated allogenic PBMCs, TM-LCL and anti-CD3 antibody. It was shown by flowcytometry that CAR expression was more than 80% in each CAR-T cell condition. 13 days after expansion, 10 Mio cells of each condition were frozen for analysis of splicing variants.

### Generation of CAR-T cells for comparability studies

CD4+ and CD8+ T cells were isolated from leukocyte reduction chamber systems by Pancoll gradient centrifugation and magnetic bead separation and activated with Transact and IL2. After two days, T cells were transfected with SB100X-encoding mRNA and CAR-encoding MC (SBorig or SBdellntr) using a Maxcyte electroporation device. T cells were expanded in T cell medium supplemented with 50 IU/ml IL-2. At day 12, T cells were harvested, counted and CAR-expression as well as T cell phenotype were analyzed by flow cytometry. Afterwards, CAR-positive CD4^+^ and CD8^+^ CAR-T cells were mixed at a 1:1 ratio and used for comparability studies.

### Comparability studies on SBorig and SBdellntr SB transposon vectors

T cells were either transfected with the SBorig or the SBdellntr MCs or left untransfected. For functional testing, they were stimulated with antigen-expressing or antigen negative target cells at different ratios. Cytotoxic/cytolytic effect was analyzed after 4 and 24 hours in a luminescence-based killing assay. After 24 hours of co-incubation, supernatant was drawn from the cells and analyzed for Interleukin-2 (IL-2) and Interferon-y (IFN-y) concentration by ELISA assay. Further, T cells were labelled with CFSE and subsequently incubated with antigen-expressing or antigen negative target cells. After 72 hours, proliferation of T cells was analyzed by measuring CFSE-dilution by flow cytometry.

---

**Supplementary Figure 1.**
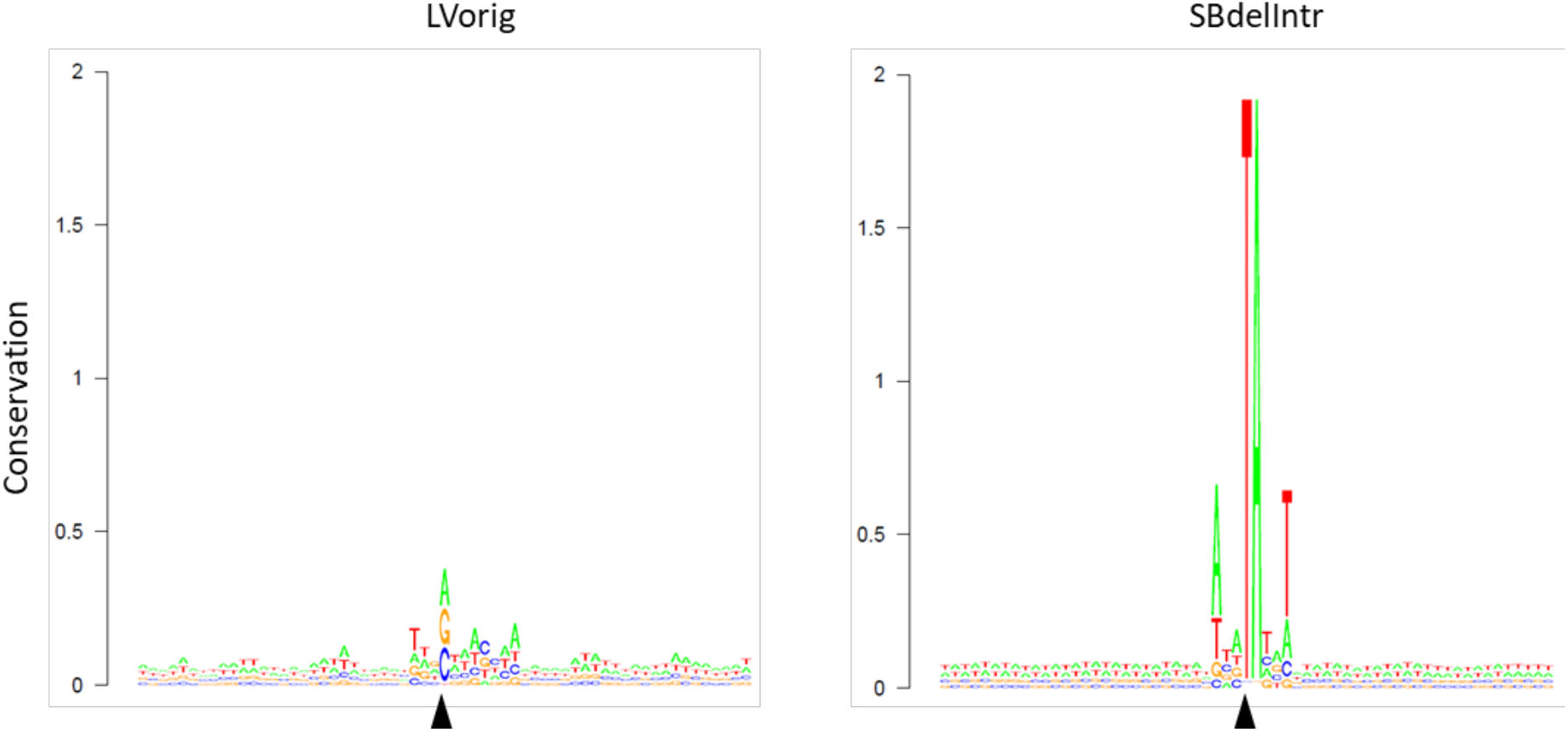
Consensus target sequences of the lentivirus and SB insertion sites in the genome of CAR-T cells. Shown are the sequence logos of the nucleotide frequencies of the genomic the insertion sites in 60 nucleotide windows around the integration positions (black arrows) of the original LV and the HTLV-1 LTR deleted version of the SB vectors. Sequence conservation (on the *y*-axis) is measured by the frequency of the nucleotides at each position (maximum conservation is log_2_ 4=2).

